# High-resolution ISR amplicon sequencing reveals personalized oral microbiome

**DOI:** 10.1101/320564

**Authors:** Chiranjit Mukherjee, Clifford J. Beall, Ann L. Griffen, Eugene J. Leys

## Abstract

**Background:** Sequencing of the 16S rRNA gene has been the standard for studying the composition of microbial communities. While it allows identification of bacteria at the level of species, it does not usually provide sufficient information to resolve at the sub-species level. Species-level resolution is not adequate for studies of transmission or stability, or for exploring subspecies variation in disease association. Current approaches using whole metagenome shotgun sequencing require very high coverage that can be cost-prohibitive and computationally challenging for diverse communities. Thus there is a need for high-resolution, yet cost-effective, high-throughput methods for characterizing microbial communities.

**Results:** Significant improvement in resolution for amplicon-based bacterial community analysis was achieved by combining amplicon sequencing of a high-diversity marker gene, the ribosomal operon ISR, with a probabilistic error modeling algorithm, DADA2. The resolving power of this new approach was compared to that of both standard and high-resolution 16S-based approaches using a set of longitudinal subgingival plaque samples. The ISR strategy achieved a 5.2-fold increase in community richness compared to reference-based 16S rRNA gene analysis, and showed 100% accuracy in predicting the correct source of a clinical sample. Individuals’ microbial communities were highly personalized, and although they exhibited some drift in membership and levels over time, that difference was always smaller than the differences between any two subjects, even after one year. The construction of an ISR database from publicly available genomic sequences allowed us to explore genomic variation *within* species, resulting in the identification of multiple variants of the ISR for most species.

**Conclusions:** The ISR approach resulted in significantly improved resolution of communities, and revealed a highly personalized, stable human oral microbiota. Multiple ISR types were observed for all species examined, demonstrating a high level of subspecies variation in the oral microbiota. The approach is high-throughput, high-resolution yet cost-effective, allowing subspecies-level community fingerprinting at a cost comparable to that of 16S rRNA gene amplicon sequencing. It will be useful for a range of applications that require high-resolution identification of organisms, including microbial tracking, community fingerprinting, and potentially for identification of virulence-associated strains.

## BACKGROUND

Short read sequencing of specific hypervariable regions of the 16S rRNA gene has been the standard for studying the composition of microbial communities for more than a decade. The technique offers several major advantages, such as high throughput, established bioinformatic pipelines and reference databases, and low per sample cost [1]. Use of this approach has revealed the remarkably diverse character of the microbiota [2] and significantly advanced our understanding of its role in human health and disease in multiple body sites [3–9]. Microbiome studies are increasingly focusing on functional relationships, and functional differences among bacterial strains of the same species have long been observed. More recent evidence from comparative genomics has shown a significant amount of variation within the genomes of different strains of the same species [10]. These genetic variations often lead to significant phenotypic differences among strains [11], which can result in varying degrees of pathogenicity [12].

While in most cases the 16S rRNA gene allows identification of bacteria at the level of species, it does not usually provide sufficient information to resolve strains within a species. Therefore, community profiling at a finer resolution than that provided by the 16S rRNA gene will be essential for better understanding the role of bacteria within systems. In addition, while the oral cavity is colonized by a diverse bacterial community [13], many species are widely shared amongst all human hosts [2]. Metagenomic studies of the gut [14–16] and skin [17] microbiomes have shown that microbial profiles show little individuality at the level of species, but high individual specificity when characterized at the subspecies level. Thus, when the operational unit of taxonomy is at best bacterial species, significant changes in microbial communities might not be detected. This is problematic for studies of bacterial transmission or stability over time.

Strain-level analyses have previously been conducted with targeted methods such as RFLP and MLST, a good example being epidemiological studies of *Mycobacterium tuberculosis* [18]. However these approaches are limited by their focus on a select set of organisms and are not suitable for community-level analysis. Another method, single nucleotide variant (SNV) analysis by whole metagenome shotgun sequencing, has recently been employed to explore strain-level variations within complex microbial communities [14, 17]. This approach allows community level characterization, and has the potential to reveal the entire genomic content of the samples [19]. However, metagenomic approaches require very deep sequencing that can be cost-prohibitive for large scale studies. Determining the relationships of individual variants among genomes, especially for highly diverse communities such as the oral microbiota, also remains challenging. Thus, there is a need for high-resolution, cost-effective, and high-throughput methods for characterizing microbial communities at the subspecies level.

An alternative region of the ribosomal operon, the intergenic spacer region (ISR) between the 16S and 23S rRNA genes provides important advantages as a target for subspecies-level community analysis. Its variability as compared to the 16S rRNA gene is sufficient to enable characterization beyond species level [20]. Sequences from the ISR have been used for detecting intra-species diversity for several bacteria such as *Listeria* spp. [21], *Lactobacillus* spp. [22], and for distinguishing viridans group streptococci [23] and diverse bacteria from clinical samples [24]. Our previous studies using heteroduplex analysis and targeted sequencing of the intergenic spacer region showed clear differential distribution of strains of *P. gingivalis* between subjects with periodontitis and periodontally healthy subjects, thus establishing a link between ISR phylogeny and disease-associated phenotype [25–29].

The ISR usually includes one or more tRNA genes, but much of it is non-coding, explaining its high variability. In addition to the high degree of variability as compared to the 16S gene that allows characterization beyond species level, the flanking conserved regions allow the use of universal primers for comprehensive community level explorations. Recently, high-throughput short read sequencing of the ISR for species-level analysis has been reported. Ruegger *et al*. showed improved resolution by short read sequencing of the ISR, as compared to 16S regions [30]. Schanche *et al*. also used high-throughput ISR sequencing and *de novo* OTU binning, suggesting microbiome transmission from mother to child [31]. However, subspecies level population structures were not explored.

A recent advance in 16S amplicon sequence analysis over OTU clustering methods has been developed by Callahan *et al*. [32]. This high-resolution amplicon sequence processing pipeline, DADA2, uses a statistical error-modeling approach to de-noise raw sequencing reads to infer true “biological variants,” and can distinguish sequences with up to single nucleotide differences. The DADA2 approach has been shown to improve resolution of 16S amplicon sequencing in several microbiome studies [8, 33–35].

We have developed a community analysis strategy combining amplicon sequencing of a high diversity marker gene, the 16-23S ISR, along with state-of-the-art bioinformatic processing software, DADA2, to provide significantly improved resolution of bacterial community composition. We show a comparison of the resolving power of this new approach to that of both standard and high-resolution 16S-based approaches. We tracked microbial communities from dental plaque of 5 adult subjects over a 1-year period. The ISR-sequencing technique combined with DADA2-based processing allowed resolution of individual species into multiple subspecies variants and revealed highly-personalized bacterial profiles for individual subjects. This efficient, high-throughput, high-resolution approach will provide greater depth for future beta-diversity based microbiome analyses and make affordable large-scale tracking of bacterial strains in comprehensive clinical studies possible.

## MATERIALS AND METHODS

### Study Design

Participants for this IRB approved study were male and female adults with a minimum of 24 remaining teeth. Exclusion criteria were antibiotic therapy or professional cleaning within the previous 3 months, requirement for antibiotic coverage before dental treatment, a chronic medical condition requiring the use of immunosuppressant medications or steroids, diabetes or HIV.

### Sample Collection

Five subjects were sampled 6 times over a period of a year. The five most anterior teeth in upper and lower right quadrants were sampled, and samples from each subject were pooled. Subgingival plaque was collected by removing excess saliva and supragingival plaque with a cotton roll and placing four sterile, medium endodontic paper points in the mesial sulcus of each tooth using aseptic technique and removing after 10 seconds. Samples were placed in 200 μL buffer ATL (Qiagen, USA) and stored at − 20°C until DNA extraction.

### Bacterial Genomic DNA Extraction

Genomic DNA was extracted from the samples using a previously described protocol [36] with minor changes to optimize yield. Briefly, thawed samples were first incubated with an additional 300 μl ATL and 40 μl Proteinase K (Qiagen, USA) at 56°C for 2 hours. After the solution was separated from the paper points by centrifugation through a perforation in the tube, it was homogenized with 0.25 g of 0.1 mm glass beads in a Mini-Beadbeater-16 (BioSpec Products, USA) for 60 seconds at 3450 rpm. Genomic DNA was purified using QIAamp DNA Mini Kit (Qiagen, USA) according to the manufacturer’s directions and eluted in 30μl of buffer AE (Qiagen, USA).

### Sequencing Library Preparation

Bacterial DNA from each sample was used to prepare two separate amplicon sequencing libraries, a 16S V1-V3 library and a 16S V8-V9-ISR library, targeting the respective regions of the ribosomal operon with locus-specific primers (**Supplementary Table S1**). Both sets of amplicon libraries were prepared based on the Illumina 16S Metagenomic Sequencing Library Preparation protocol (Illumina, USA), a two-step protocol in which the region of interest is first amplified with gene-specific primers and then dual indices are added through a brief second PCR. Optimizations were made to the Illumina protocol to allow for high-fidelity amplification and automation. For each sample, purified DNA was adjusted to 5 ng/μL concentration and 2 μl of DNA was PCR amplified in a total volume of 25 with Accuprime Taq DNA Polymerase, High Fidelity (ThermoFisher Scientific) using primers (IDT, USA) composed of region-specific sequences (Table S1) and attached Illumina adapter sequences. Forward Illumina adapter used was TCGTCGGCAGCGTCAGATGTGTATAAGAGACAG and reverse Illumina Adapter used was GTCTCGTGGGCTCGGAGATGTGTATAAGAGACAG. PCR parameters consisted of an initial denaturation step at 94°C for 2 minutes, followed by 25 cycles of denaturation at 94°C for 30 secs, annealing at 55 °C for 30 seconds and extension at 68 °C for 1 min followed by a final extension at 72°C for 5 min. PCR products were purified with the Agencourt AMPure XP PCR Purification system (Beckman Coulter, USA) using manufacturer’s guidelines. This was followed by a subsequent 8-cycle PCR amplification step (temperature and timing as before) to add dual indexing barcodes for multiplexing, with barcode sequences derived from the study by Kozich *et al*. [37]. The index PCR products were further purified using the AMPure system as above and quantified using Quant-iT™ High-Sensitivity dsDNA Assay Kit (Invitrogen, USA) on a Spectramax Microplate reader (Molecular Devices, USA). Sequencing library validation was performed with the Fragment Analyzer (Advanced Analytical, USA) using the dsDNA 915 Reagent Kit and the manufacturer’s protocol. The validated libraries were adjusted to 10 nM concentration and pooled together. All pipetting steps were performed using a Biomek 4000 Liquid Handling Automation Workstation (Beckman Coulter) for consistency. Both the 16S V1-V3 and 16S V8-V9-ISR libraries were sequenced on the Illumina MiSeq Platform with 300 base pair paired-end chemistry.

### Human Oral ISR Database

A list of the most abundant and prevalent dental plaque-associated oral bacteria was generated by community profiling based on 16S V1-V3 results. Available genomes for these bacteria were downloaded from NCBI’s GenBank [38] database. For finished genomes, a custom R script implementing command line RNAmmer [39] (version 1.2) was used to identify 16S and 23S rRNA genes and extract the 16-23S internal spacer region (ISR) sequences. For draft genomes, another custom R script utilizing command line BLASTN [40] application 2.5.0 with ISR-flanking primers was used to identify and extract ISR sequences. When creating our 16S database of oral bacteria, CORE, we could not distinguish between certain closely related species and therefore found it necessary to group them together. Distinct phylogenetic clades were observed within each of these species groups. However, we were not able to assign each clade to a specific species. As a result, to avoid ambiguous taxonomic assignments, and maintain consistency with our 16S database, we combined ISRs from these species into groups matching those in our CORE database. A tabular representation of this database in included in Supplementary Table S2.

### Comparison of Diversity between 16S and ISR

Genomes of 14 different species from the genus *Streptococcus*, one of the most abundant bacteria in the dental plaque communities, were downloaded from NCBI GenBank, and the 16S V1-V3 region and 16-23S ISR were extracted *in silico* using locus-specific primers and custom R scripts. The 16S and ISR sequences were aligned independently using MUSCLE multiple sequence aligner [41] implemented within Unipro UGENE bioinformatics toolkit (**Figure 1**) [42].

**Figure 1.**
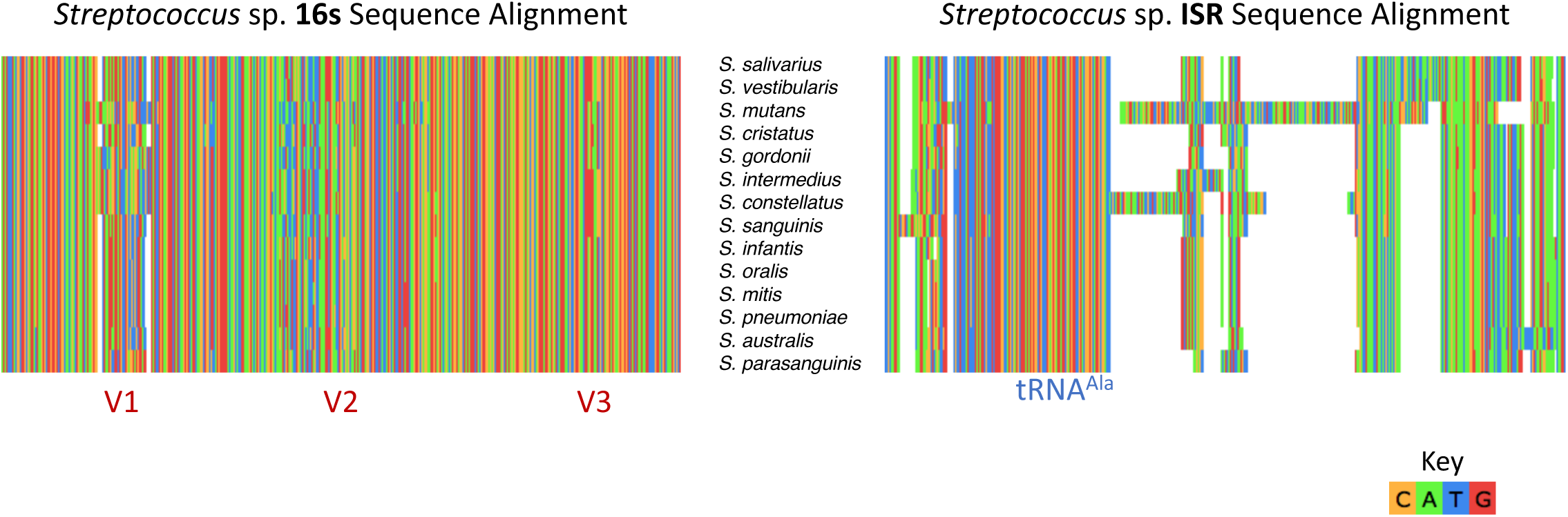
Higher diversity in the 16-23S Intergenic Spacer Region (ISR) compared to the 16S V1-V3 among oral streptococci. **Left:** Alignment of the 16S V1-V3 regions of common oral *Streptococcus* sp. shows alternating conserved and hypervariable regions. The limited amount of variation makes it difficult to distinguish between oral streptococci using 16S rRNA gene sequencing. **Right:** Alignment of the ISR sequence of the same *Streptococcus* sp. show much higher variation, as evident from large gap regions, with the most prominent conserved region being the alanine tRNA gene.

### Bioinformatic Processing of Sequencing Data

#### 16S V1-V3 Paired Sequences

De-multiplexed, paired reads from the 16S V1-V3 library were processed using an in-house bioinformatic pipeline (shell script) for analyzing 16S amplicon sequencing data. Forward/reverse read pairs were combined into contiguous sequences using Mothur [43]. Unexpected length sequences and those with >10 ambiguous bases were filtered out and PCR primer sequences were removed. The resulting sequences were then filtered to keep only those reads which had average Phred quality scores ≥ 28, using a script based on BioPython [44] 1.63 (Python 2.7.6). Next, these high quality reads were aligned with the command line BLASTN application 2.5.0 against the OSU CORE reference database of human oral 16S sequences [45]. A custom PHP script was used to recalculate percent identities (avoiding counting ambiguous bases as mismatches) and to filter by the length of the alignments. Sequences which matched ≥ 98% and had a minimum alignment length of 400 base pairs with the reference database entries were classified taxonomically as the highest identity match. The blast results were then tabulated to generate a sample vs. species-OTU table, which was used for further analysis.

#### 16S V8-V9 Sequences

Reads consisting of 16S V8-V9 sequences, which were sequenced as Read2 in the 16S V8-V9-ISR amplicon sequencing procedure, were processed using our standard 16S amplicon sequence processing pipeline as described above, modified for unpaired reads. The quality trimming parameters used for these sequences were an average score of ≥ 25 over a window size of 30 bases. High quality sequences which passed this screen were assigned species-level taxonomy using command line BLAST against OSU CORE database as described above, but with a minimum alignment length of 200 base pairs. This enabled us to compare the species-OTU level diversity between the V1-V3 and V8-V9-ISR primers.

#### ISR Sequences

The ISR reads, sequenced as read1 in the 16S-V8-V9-ISR amplicon sequencing procedure, were processed using the Bioconductor package *dada2* (version 1.6) to infer exact sequence variants, with modifications for single reads. The raw FASTQ files were filtered with *dada2* package function fastqFilter as per the following parameters: the sequences were truncated to a length of 295 bases (truncLen=295), the PCR primers were trimmed off (trimLeft=15), and reads with more than two expected errors (maxEE=2) were discarded. The sequencing error rates were estimated *(dada2* function learnErrors) and the filtered sequences were then *dereplicated (dada2* function derepFastq) to generate unique sequences. These unique sequences were then processed using the DADA2 algorithm *(dada2* function dada) to infer exact amplicon sequence variants (ASVs). Chimeric sequences were removed *(dada2* function removeBimeraDenovo) and a sample vs ISR ASV table analogous to OTU-tables was generated (phyloseq function otu_table) [46], which was used for further community-level distance-based analysis.

#### 16S V1-V2 Sequences

To evaluate the effect of the bioinformatic pipeline itself, we also processed the unpaired read1s from the 16S V1-V3 data with DADA2 to generate high-resolution 16S amplicon sequence variants, using parameters similar to those used for the ISR sequences. The resulting sample vs 16S ASV table was used for distance analysis.

#### Data Analysis & Visualization

**Distance-based community analysis:** Community composition tables generated from the 3 different pipelines were rarefied using the R Function *Rarefy* (R package: GUniFrac). Pipeline statistics were generated based on rarefied counts data. The rarefied counts data for each pipeline were transformed to presence/absence tables, and these tables were used to calculate the number of subject-specific and unique ASVs. A box-and-whisker plot of centroid distances was evaluated from multivariate homogeneity of group dispersions, computed using the R function *betadisper* (package: vegan). Wilcoxon Rank Sum test was performed using R function *wilcox.test* (package: stats). Nonmetric multidimensional scaling (NMDS) plots were generated with the R function *metaMDS* (package: vegan), based on Bray-Curtis dissimilarities, from the presence/absence tables. Ellipses were drawn around each subject group using the R function *ordiellipse* (package: vegan) at 95% confidence intervals. Hierarchical clustering and calculation of p-values was performed using R package *pvclust* (package: pvclust), using the “correlation” distance measure, and the agglomerative clustering method “complete”, with 1000 bootstrap replicates. Clusters with bootstrap probability ≥ 95% were highlighted with R function *pvrect* (package: pvclust). Stability plots were constructed by plotting Bray-Curtis distances between initial and successive time point samples for each subject. Slope was calculated by fitting a linear mixed-effects model using R function *lm* (package: stat). Random forest classifier model training was performed using the R function *randomForest* (package: randomForest). Accuracy statistics were obtained from the R function *confusionMatrix* (package: caret).

**Species-specific ISR-type analysis:** ISR ASVs generated using DADA2 from the ISR-amplicons were mapped against the Human Oral ISR Database using custom BASH scripts implementing command line BLASTN application and PHP script as before. The BLAST results were processed using R. For BLAST alignments over 90% sequence identity no conflicts were observed in species assignment, and therefore this threshold was selected as a cutoff for assignment of taxonomy. All amplicon sequence variants (ASVs) that matched with the ISR-database sequence of species at ≥ 90% identity and were present in more than 1 sample were thus marked as the ISR-types of that species. This allowed us to resolve most common oral bacteria into a number of ISR-type strains. ASVs that were not identified by BLAST against the ISR database were clustered using USEARCH v8 [47]. Representative sequences from each cluster were mapped against NCBI’s GenBank database [48], but no additional species were identified.

For ISR-type population structure analysis, sample counts were standardized for sequencing depth using the R function *decostand* (package: vegan). Distribution of ISR-types were plotted using the R function *heatmap*.2 (package: gplots). Rarefaction curves were plotted using R function *rarecurve* (package: vegan).

## RESULTS

### Higher diversity in the ISR compared to 16S V1-V3

We evaluated the potential of the ribosomal 16-23S ISR for exploring diversity within species by determining how much variation was present among oral bacteria within the ISR compared to the 16S V1-V3 hypervariable region. Oral streptococci are among the most difficult organisms to distinguish by 16S-based analysis. Multiple sequence alignments for the two regions for 14 species of streptococci (**Fig. 1**) showed substantially higher diversity among the ISR sequences, with the main region of homology being the locus coding for an alanine tRNA. Similar patterns were observed for species from other genera (data not shown). These alignments indicated, as expected, the ISR could be targeted as a higher resolution marker for amplicon sequencing of the oral microbiota.

### Clinical results for evaluating ISR-based amplicon sequencing

To evaluate the effectiveness of an amplicon sequencing strategy based on targeting the ISR, subgingival plaque from five subjects (S1-5) was sampled six times (T1-6) over a period of 11 months at intervals ranging from 0.9 to 3.5 months (**Fig. 2**). Each time the entire group was sampled on the same day, and the same teeth were sampled. Four subjects were periodontally healthy (PD ≤ 3mm at all sites). Subject #5 had PD ≥ 5mm at one site, and the sampling strategy was slightly altered for this subject: first molars with PD ≥ 5mm were sampled in lieu of the 2^nd^ bicuspid. Bacterial genomic DNA was extracted and two amplicon sequencing libraries were prepared for each sample, targeting the 16S V1-V3 hypervariable region and the 16S V8-V9-ISR, using appropriate locus-specific amplification primers (**Fig. 2, Table S1**).

**Figure 2.**
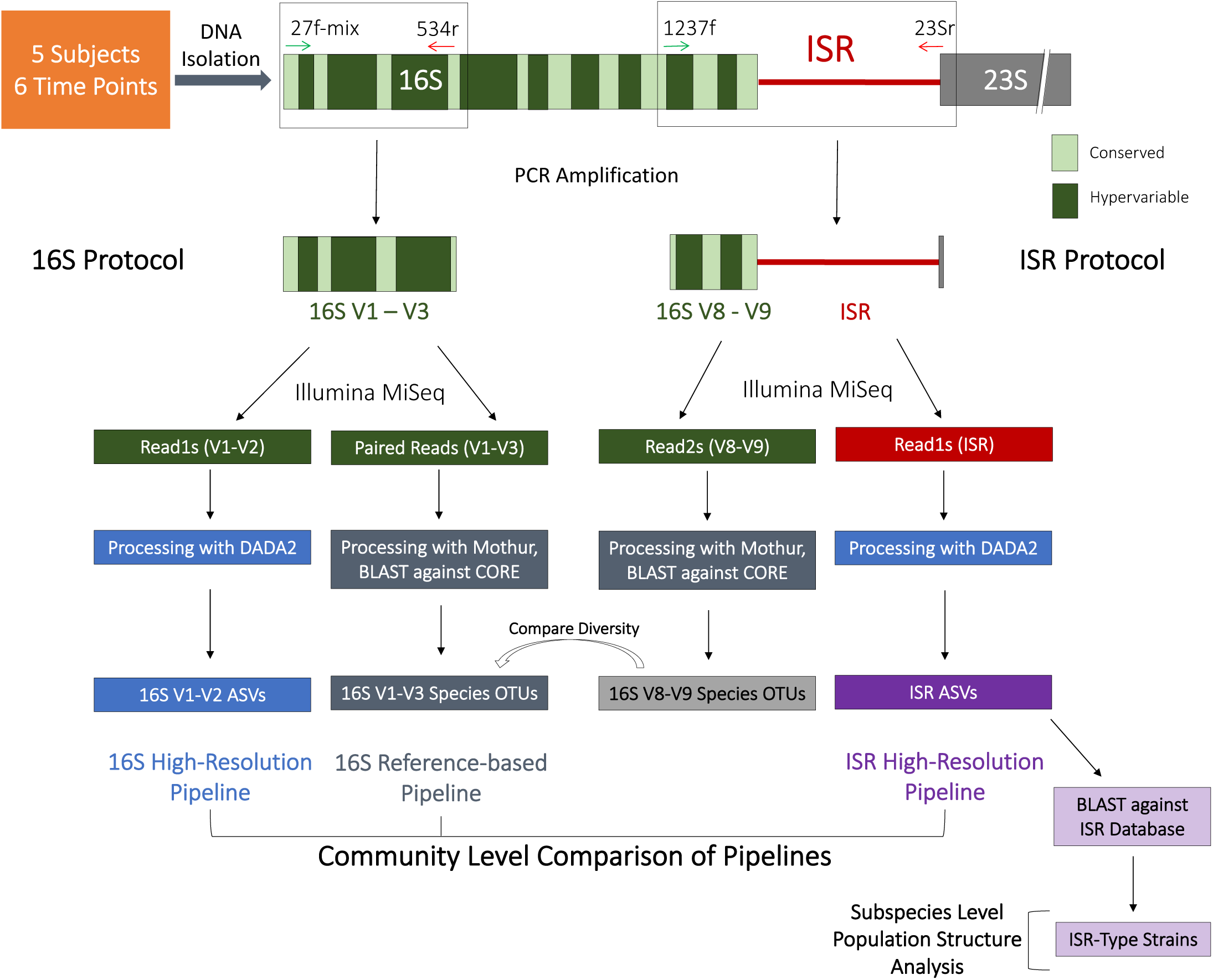
Schematic of the PCR amplification, sequencing and bioinformatic processing pipeline for this study. The 16S V1-V3 and 16S V8-V9-ISR regions were amplified separately, with validated universal primers. Amplicons were sequenced using Illumina MiSeq with 2×300 bp paired-end chemistry. Sequence reads were processed in 3 different bioinformatic pipelines as shown, and community composition was compared to evaluate performance of each pipeline. The ISR reads were also used for subspecies level population structure analysis.

### ISR pipeline resolved microbial communities with greatest resolution

Sequencing reads generated from the 16S V1-V3 hypervariable region and the 16S V8-V9-ISR libraries were processed in the following ways:

- **16S Reference-based analysis pipeline:** Paired reads from the 16S V1-V3 amplicon library were merged, quality filtered and aligned with reference database OSU CORE [45] to assign taxonomy at the species level
- **16S High-Resolution analysis pipeline:** Single reads from the 16S V1-V3 amplicon library containing the V1-V2 region were processed using the high-resolution sample inference pipeline DADA2 [32].
- **ISR High-Resolution analysis pipeline:** Single reads from the 16S-V8-V9-ISR amplicon library (containing the ISR) were also processed using the DADA2 pipeline. In addition, single reads containing the 16S V8-V9 region were processed using OSU core database as described for the 16S reference-based analysis pipeline.

Comparison of the 3 pipelines for the dataset of 30 samples is shown in **Fig. 3**. Sequences processed by the 16S reference-based analysis pipeline mapped to 359 species-level OTUs. In contrast, the same sequences processed with the 16S high-resolution analysis pipeline were binned into 1725 amplicon sequence variants (ASVs). The greatest number of variants were inferred with the ISR high-resolution analysis pipeline, with 1839 subspecies-level ASVs **(Fig. 3A)**. The simultaneous sequencing of the 16S V8-V9 region allowed us to map them to 224 species. Since 359 species were represented in the standard 16S pipeline, this indicated that the 16S-V8-V9-ISR amplicon library was not comprehensive for all species. We attribute this to region-specific primer biases and longer amplicon lengths for some species.

**Figure 3.**
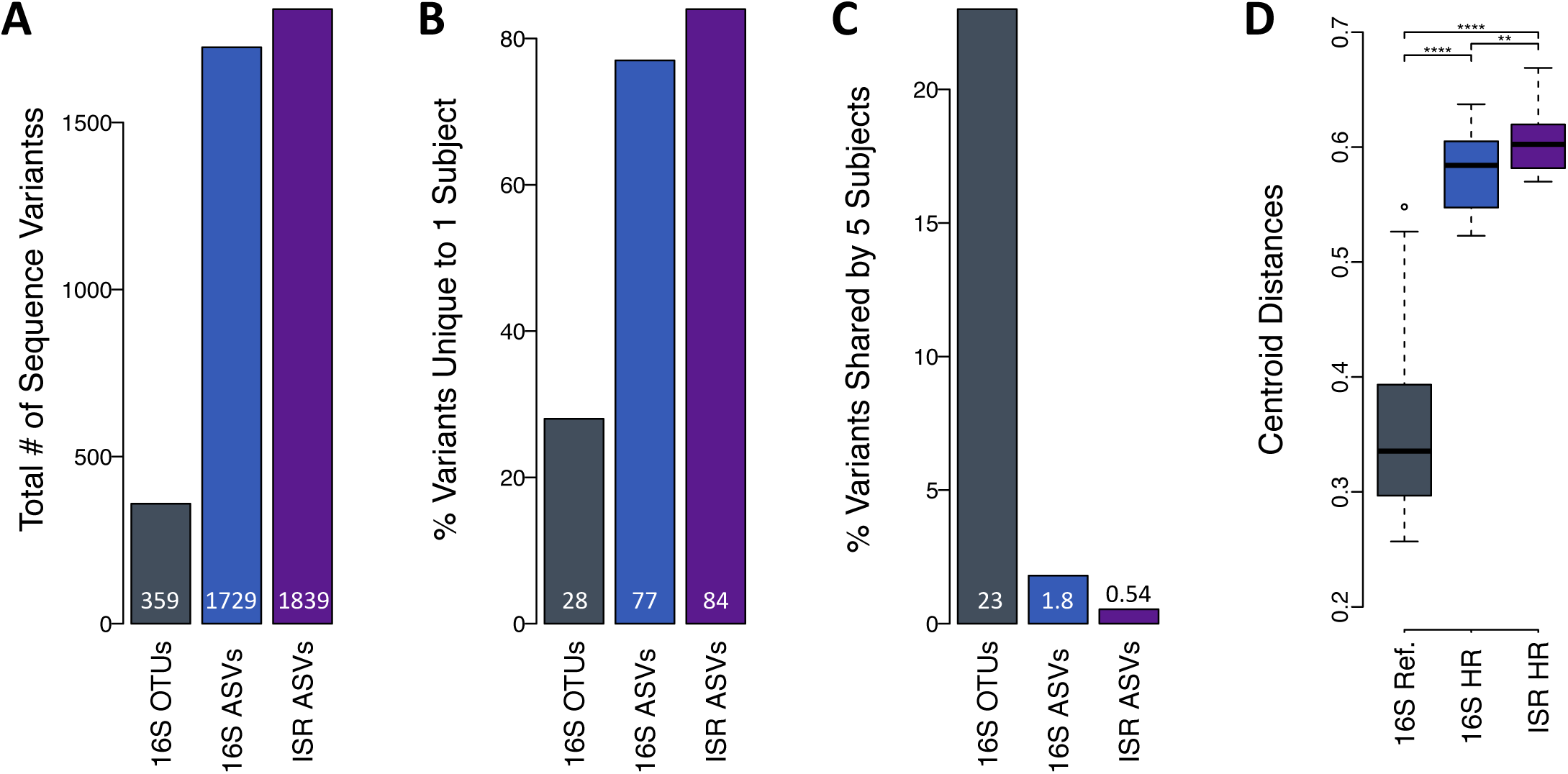
ISR pipeline provided greatest resolution of microbial communities. **A)** This panel shows the number of sequence variants generated by each of the 3 pipelines. **B)** This panel shows the percentage of variants unique to a single subject, as determined by each of the 3 pipelines. **C)** This panel shows the percentage of variants shared by all 5 subjects, as determined by each of the 3 pipelines. **D)** Box-and-whisker plot show the degree of separation between the samples, or sample resolution, as measured by calculating the Euclidean distances in principal coordinate space between the samples and the centroid, based on Bray-Curtis dissimilarities. Asterisks represent level of significance based on p-values from the Wilcoxon Rank Sum Test.

The percentage of variants unique to a single subject **(Fig. 3B)** was highest using the ISR pipeline, intermediate using the 16S high-resolution pipeline, and lowest using the 16S reference-based pipeline (84%, 77%, 28% respectively). The percentage of variants shared among all subjects **(Figures 3C)** was lowest using the ISR pipeline, intermediate using the 16S high-resolution pipeline, and highest using the 16S reference-based pipeline (0.54%, 1.8%, 23% respectively).

Community membership tables for each of these 3 approaches were used in beta-diversity analyses using Bray-Curtis dissimilarity indices to compute distances between samples and compare the resolving power of the three pipelines (**Fig. 3D**). The resolution was greatest for the ISR pipeline but both the high-resolution approaches were significantly better than the 16S reference-based pipeline.

### High-resolution ISR sequencing revealed personalized microbiome

Longitudinal samples were analyzed to determine the relationship of microbial profiles for the 5 individuals over the 1 year period. Non-metric multidimensional scaling (NMDS) plots of Bray-Curtis dissimilarity, hierarchical clustering analyses and random forest classification were performed, all based on community membership (presence/absence).

In NMDS ordinations constructed from the ISR pipeline **(Fig. 4 - Top panel right)**, all points were contained within the 95% confidence interval ellipses drawn around the centroid of each group, and there was no overlap among groups. For the 16S high-resolution pipeline, there was considerable overlap between the confidence interval ellipses **(Fig. 4 - Top panel middle)**, while the overlap for the species level 16S reference-based pipeline was extensive **(Fig. 4 - Top panel left)**. For both the 16S pipelines, at no confidence level were all the samples contained within their designated confidence interval.

**Figure 4.**
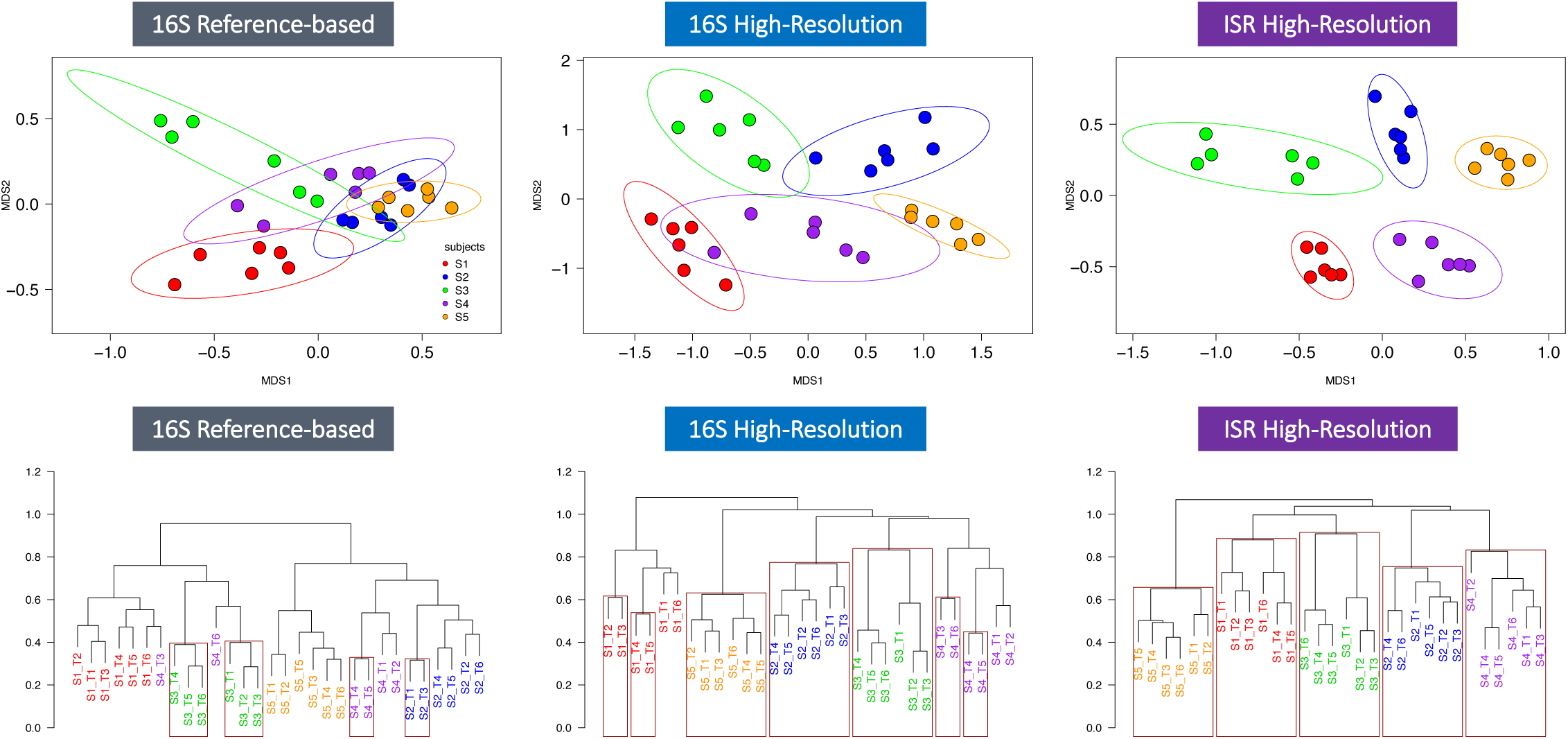
Distance-based community analysis of the high-resolution ISR pipeline data revealed personalized microbial profiles for all subjects. **Top panel**: From left to right, non-metric multidimensional scaling (NMDS) plots from Bray-Curtis dissimilarity matrices based on community membership are shown for the 3 approaches: 16S reference-database, 16S high-resolution and ISR high-resolution. Six time points are shown for 5 subjects with 95% confidence intervals. **Bottom panel:** From left to right, hierarchical clustering dendograms based on correlational distance are shown for the 3 approaches. Rectangles are drawn marking largest clades with ≥ 95% bootstrap probability.

Agglomerative hierarchical clustering dendograms based on correlational distance for the 3 pipelines were also constructed, and clades with ≥ 95% bootstrap probability were marked. For the ISR pipeline, classification accuracy was 100%, and all 5 subjects were grouped in distinct, individual clades (**Fig. 4 - Bottom panel right**). For the 16S high-resolution pipeline, classification was also 100% accurate, but bootstrap probabilities were ≥ 95% for only 3 subject clades (**Fig. 4 - Bottom panel middle**). For the 16S reference-database pipeline, none of the subject clades reached ≥ 95% bootstrap probability values, and accuracy in distinguishing individuals was poor (**Fig. 4 - Bottom panel left**).

Using a random forest classifier, the accuracy was calculated for all three pipelines by training the model on random subsamples of the community membership matrix, leaving out one sample each time and testing classification accuracy for the remaining sample. Mean classification accuracy for the ISR pipeline, 16S high-resolution pipeline and 16S reference-based pipeline were 100%, 96.7%, and 93.3% respectively.

### Stability of the oral microbiota over 1 year

We estimated the stability of the subjects’ oral microbial communities by comparing intra-subject changes over time and mean inter-subject differences for microbial profiles generated using all three pipelines. For each subject, we plotted the Bray-Curtis dissimilarity based on community membership between the baseline sample and samples collected at succeeding time points **(Fig. 5)**.

**Figure 5.**
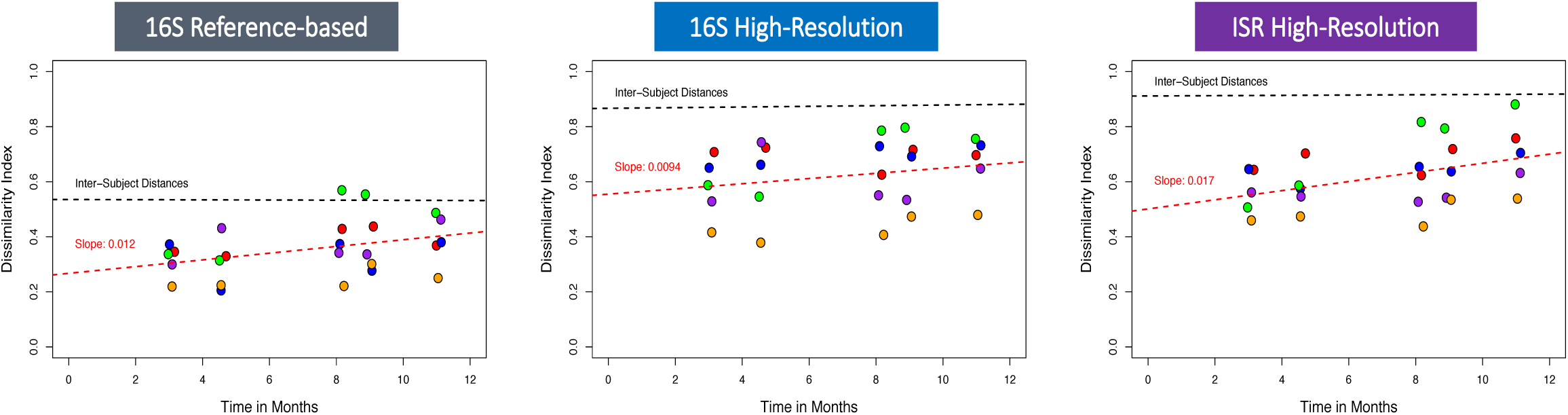
Stability of the oral microbiota over 1 year. Bray-Curtis dissimilarities between the baseline and subsequent time points for each subject are shown for the 16S reference-database (**left**), 16S high-resolution (**center**) and ISR high-resolution (**right**) approaches. The red dotted line shows the fit of the linear model for within-subject distances. For comparison, the fit for between-subject distances is shown by the black dotted line.

A low level of drift from baseline microbial profiles for subjects over time was consistently observed with all three of our approaches. However, with both the ISR and 16S high-resolution pipelines, differences within subjects over one year were always less than the mean inter-subject differences **(Fig. 5)**. With the lower resolution species-level 16S reference-based pipeline, intra-subject differences occasionally exceeded inter-subject differences.

### Creation of Oral ISR Database Enabled Species Assignment to ISR ASVs

To explore genomic variation *within* each species, taxonomic assignment of the ISR sequences at the species level is required. To accomplish this, we developed an ISR database by *in silico* extraction of the 16S-23S intergenic spacer region from over 200 publicly available genomic sequences of bacteria that were most abundant or prevalent in the 16S datasets. Currently this database contains 176 unique sequences from 60 species groups matching the CORE 16S database.

ISR ASVs which were present in more than one sample were considered for species assignment. 338 ASVs, representing 73.1% of total sequences being considered, were mapped to 31 species groups with this approach. Using this subset of ISR ASVs in distance-based analysis, we were able to resolve individuals as well as the full dataset that included the unmapped ASVs (**Supplementary Fig. S1**).

### Population structures and persistence of ISR-types

Mapping the ISR ASVs against our oral ISR database resulted in the identification of multiple genotypic variants of the ISR, or ISR-type strains, for most species. The number of ISR-types for the 15 most abundant species (with >5000 sequences each) in our sample set ranged from 5 to 41, with an average of 15.1 ISR-types per species group (**Table 1**). We selected the three species which showed maximum ISR-type diversity, *Haemophilus parainfluenzae, Granulicatella adiacens*, and the *Streptococcus mitis* group, for an in-depth analysis. These species could be resolved into 41, 38 and 33 ISR-types respectively, and were labelled Hp 1-41, Ga 1-38, and Sm 1-33, respectively, based on decreasing overall sequence abundance. Rarefaction analysis for each of the 15 species showed that generally our sequencing strategy provided sufficient depth to profile all real ISR variants for these species. Rarefaction curves for the top 3 most diverse species are shown in Supplementary **Fig. S2**.

The population structure of *H. parainfluenzae, G. adiacens*, and S. *mitis* group were similar, with a few ISR-types showing high abundance, and a large number of ISR-types present in relatively lower abundance (**Fig 6A)**. We also looked at prevalence of ISR-types among subjects (**Fig 6 B & C)**. The distribution ranged from presence in only a single subject (16/41 ISR-types for *H. parainfluenzae*, 23/39 ISR-types for *G. adiacens* and 16/33 ISR-types for S. *mitis* group) to one ISR-type of S. *mitis* group that was ubiquitous. The relative levels over time of the major ISR-types of *H. parainfluenzae, G. adiacens* and the S. *mitis* group are shown in **Figure 7**.

**Figure 6.**
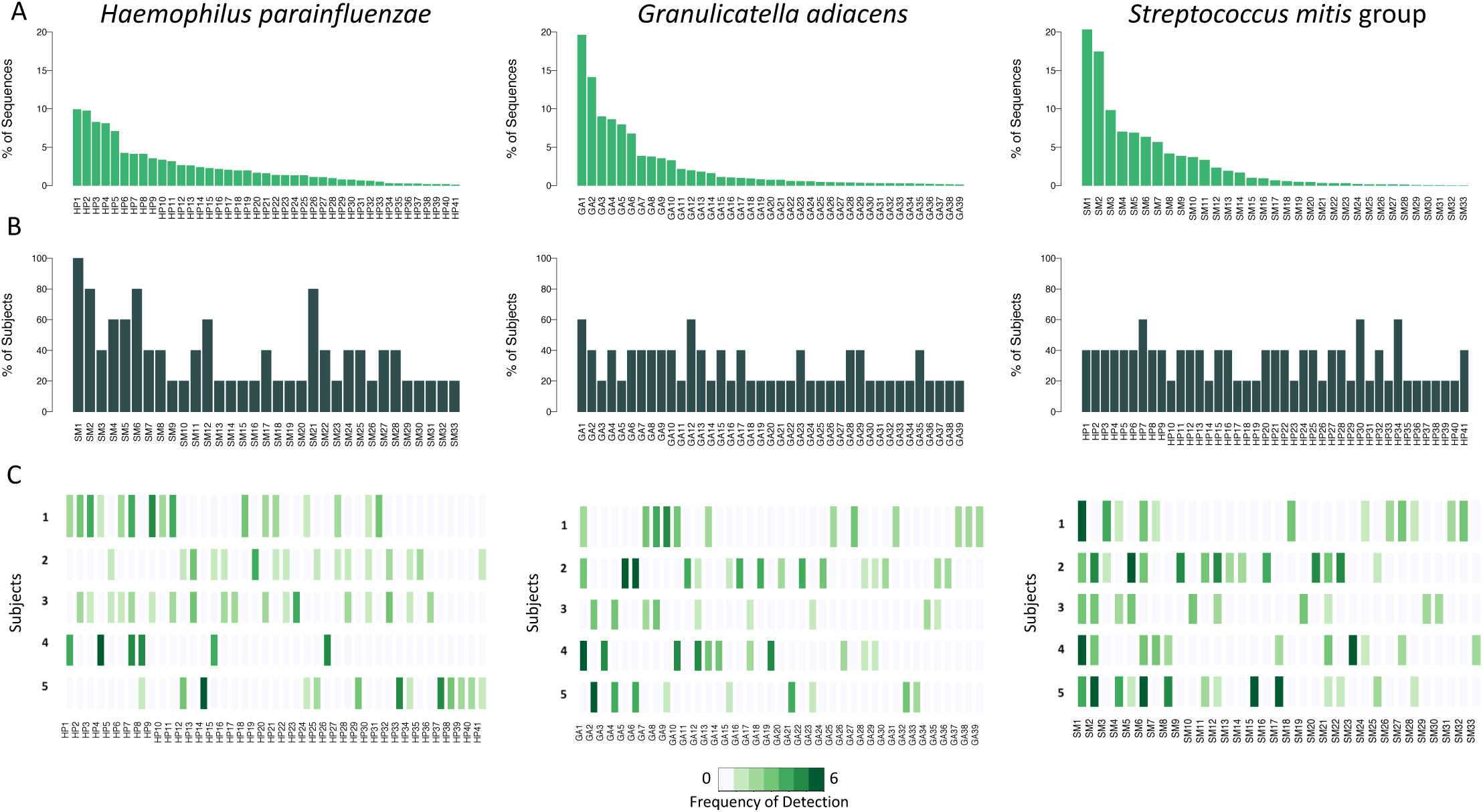
Population structure for three abundant oral bacteria. A) For each species, bar plot of relative abundance for the entire dataset is shown. The ISR-types are ranked and labelled in order of decreasing abundance, and the same order is used in the lower panels. B) Prevalence among subjects, determined by presence at any time point, is shown as bar graphs. C) Frequency of detection of ISR-types over time within subjects is shown using a heat map.

**Figure 7.**
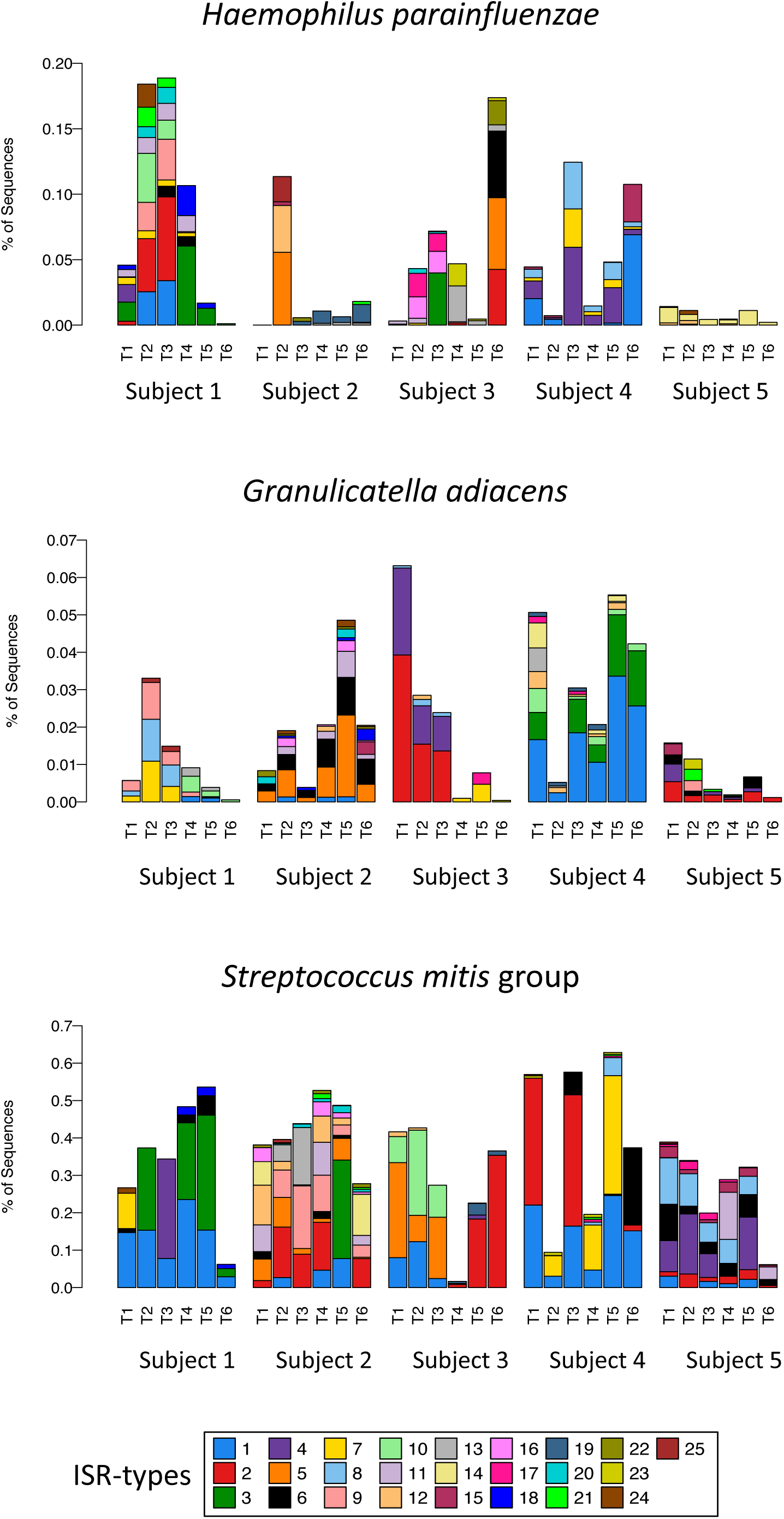
Relative abundance of ISR-Types fluctuate over time. Relative abundance of ISR-types for *H. parainfluenzae, Granulicatella adiacens* and the *Streptococcus mitis* group, for each subject, are shown in stacked bar plots. ISR-types were numbered as in Figure 6, based on their overall abundance, and only the 25 most abundant are shown. The same color scheme based on abundance was used for all 3 species.

## DISCUSSION

### Comparison of the 3 amplicon sequencing processing pipelines

Performance validation of new sequencing and processing strategies is usually done using relatively low complexity synthetic mock communities. However, the goal of this study was to explore variation at the subspecies level. Since it would not be possible to construct a mock community using laboratory strains that accurately mimics the degree of strain variation found in natural communities, clinical samples were used for comparison of pipelines. Longitudinal dental plaque samples were collected from unrelated healthy adult subjects. This study design provided a model system for measuring community stability and inter-individual variation.

Our novel approach included two improvements over standard 16S amplicon pipelines, a more variable target sequence, the ISR, and incorporation of a high-resolution probabilistic error modeling pipeline, DADA2. Oral microbial communities have been well characterized at the species level and well curated databases are available [45, 49]. Mapping 16S reads against databases has become the standard for community analysis. Without an existing comprehensive ISR database, standard reference-based processing was not possible for the ISR reads. Instead, we processed the ISR sequences with the DADA2 pipeline, and compared the results to standard 16S rRNA gene V1-V3 region amplicon sequencing processed by reference database matching, and to the same 16S sequences processed using DADA2.

Our goal was to resolve complex microbial communities into the highest possible number of statistically verified true biological variants. As expected, the highest number of sequence variants were inferred from the ISR sequences, with a 5.2-fold increase over species-level taxons. Processing the 16S reads using the DADA2 pipeline also resulted in improvement as compared to the reference-based pipeline, showing a 4.8-fold increase. The relatively small difference between these two approaches was the result of the V8-V9-ISR library being less comprehensive than the V1-V3 library, as discussed below. This suggests that future efforts focused on improving the comprehensiveness of ISR amplification protocols could improve resolving power even more.

Overall, the ISR pipeline discriminated among hosts with the greatest confidence and accuracy, and provided the highest resolution for amplicon sequencing-based characterization of bacterial communities. Individuals could be perfectly resolved, with no overlap seen among samples from different subjects in NMDS ordinations and hierarchical clustering dendograms. The 16S highresolution pipeline did not perform as well at the ISR pipeline, but it did perform better than the 16S reference-based pipeline.

### Personalized Oral Microbiota

This year-long study of 5 individuals provides the strongest evidence to date that oral microbial communities are highly personalized, as demonstrated by clustering and machine learning based approaches. The ISR pipeline showed 100% accuracy in predicting the correct source of an oral clinical sample. It is interesting that the subjects were all in near daily contact with each other, although none lived together or were related, suggesting that casual contact is not an important determinant of microbial transmission. We had also collected plaque samples from additional subjects, though not for all the time points. Including these samples in the distance based analysis did not alter the distinctiveness of each subject’s microbiome (data not shown).

Personalization of the oral microbiota has been observed in at least two previous, similar studies, although neither were able to perfectly resolve individuals. In a re-analysis of a 16S longitudinal supragingival plaque dataset, using an “oligotyping” approach based on individual nucleotide positions (Utter et al., 2016), resolution was comparable to that seen in the current study for 16S-DADA2, but less than the ISR-DADA2. In both the Utter study and another study using 16S sequencing and OTU clustering [50], the accuracy observed for reference based mapping of 16S was much lower, similar to that observed for 16S reference-based approach in the present study.

### Microbial Stability

In addition to examining differences among subjects as described above, stability over time for each subject was evaluated. Using the ISR approach, for each subject oral microbial community composition at baseline was compared to that observed at each succeeding time point **(Fig. 5)**. Although individuals’ microbial communities exhibited slight drift in membership and levels (data not shown) over time, that difference was always smaller than the differences between any two subjects, even after one year. In contrast to results using the ISR, with a conventional 16S reference mapping-based pipeline **(Fig. 5)**, intra-subject differences occasionally exceeded inter-subject differences, showing that fine-scale techniques give a more accurate window on community stability.

Our observation of the change in microbial profiles over time are in agreement with the latest findings from the Human Microbiome Project, where a baseline level of intra-individual variation was observed by analyzing strain level communities from metagenomic sequencing data [19]. Overall, our findings with regard to the stability and individuality of the plaque microbiome are similar to those seen in other sites, including tongue and saliva microbiome [51, 52], skin [17] and gut microbiome [14, 15], and recently in fecal microbiome [16]. This provides further support for the utility of subspecies level microbial community characterization as a method for ‘microbial fingerprinting’ as has been previously suggested [53].

### Exploring subspecies-level population structure & dynamics

The ISR locus provided sufficient diversity to allow exploration of individual population structures for many of the species found in the samples. Species level taxonomy was assigned to the ISR ASVs using the ISR database described above.

Development of this database will be an ongoing project as new genomes are sequenced. To avoid inclusion of potential amplification artifacts, ASVs were only considered if they were detected in at least two samples, even though this may have resulted in underestimation of population diversity.

Most of the common oral species exhibited considerable ISR-type heterogeneity, demonstrating a high level of subspecies variation in the oral microbiota. *H. parainfluenzae* demonstrated the most, with 41 ISR-types. This is consistent with a recent metagenomic strain profiling showing that this bacterium has high strain diversity compared to other human associated microbes [19]. Rarefaction analysis showed that sequencing depth was sufficient to estimate population structures for all 15 species we analyzed. Three species were selected for more detailed profiling based on highest diversity and overall abundance, *Haemophilus parainfluenzae, Granulicatella adiacens*, and *Streptococcus mitis* **(Fig 6)**.

The degree to which ISR-types were shared amongst subjects, or prevalence, ranged from one S. *mitis* group variant that was shared by all subjects, to a large number of variants from each species that were present in only one subject (**Fig 6B & C**). These individually unique variants are major contributors to the ability to discriminate among subjects based on beta-diversity analysis (data not shown). Surprisingly, abundance and prevalence were not well correlated; some of the less abundant ISR-types were found in multiple subjects **(Fig 6 A & B)**.

Population dynamics for 3 species are shown for 5 subjects in Figure 7. Many ISR-types were retained over one year, but considerable fluctuation in levels and frequent loss and gain of ISR-types were observed (Fig 7).

### Relationship between the ISR and rest of the genome

While our results show that the ISR technique provides a powerful tool for distance-based beta diversity analysis and tracking variants within bacteria species, the functional significance of ISR-type variability has not yet been established. The relationship between the ISR and the rest of the genome can vary from species to species. One study involving several species of phototrophic bacteria belonging to the order *Rhizobiales* showed that 70% DNA hybridization was associated with 92% ISR sequence similarity [54]. How closely this correlates with conventional phenotype-based strain designations is not presently fully understood. Our future work in this area will focus on establishing genomic and functional correlations for ISR-types.

Certain bacterial species also possess multiple ISR types, which may vary in length and the inclusion of tRNA genes. Although bacterial genomes usually contain multiple copies of the ribosomal operon, for many of the published genomes we analyzed, these multiple copies of the ISR were identical. However, some taxa such as Veillonella spp., *Aggregatibacter actinomycetemcomitans* and *Porphyromonas gingivalis* have multiple non-identical ISR sequences. This information is currently available for a limited number of species and strains. Potential methods for detecting non-identical ISRs might include identifying integer multiples or significant concordance of ISR-types within samples.

### Comparison between the 16S V1-V3 and 16S V8-V9-ISR libraries

A portion of the 3’-end of the 16S rRNA gene was included in our original ISR amplicon, with the intention of allowing direct mapping of ISR variants to species-level taxa using the V8-V9 region, but the following inherent constraints limited its utility. In Illumina paired-end sequencing, the second sequencing reads are lower quality compared to the first reads. Since the ISR read quality was critical, these were sequenced first, followed by the 16S V8-V9 reads. This resulted in the loss of over 25% of the V8-V9 reads during quality and length filtering. Additionally, species assignments using the V8-V9 region are not as accurate as those based on the well validated, standard V1-V3 sequences [55]. So we instead developed the ISR database to map the ISR variants to species level taxa, and relied on separate amplification of the V1-V3 region for species level community characterization.

Although it was not ideal for species mapping, the inclusion of the V8-V9 region in the ISR fragment did allow comparison of microbial diversity of the 16S-ISR library with the standard, comprehensive 16S-only library. Mapping the V8-V9 sequences generated a subset of these species-level taxons (222 of the 359) identified from the V1-V3 region in our dataset. Comparison of the species taxons in the two groups showed that many of the species missing in the V8-V9 sequences had longer (>500 bp) ISRs, resulting in very long amplicons (>800 bp), which do not amplify or sequence as well as the shorter amplicons from species with smaller ISR regions. Since it is possible to assign taxonomy without the V8-V9 region, sequencing just the shorter 16-23S ISR fragment may allow greater diversity in the ISR libraries due to smaller amplicon lengths.

### Application of the ISR technique

High-resolution community characterization, beyond species level, is currently achievable only by whole metagenome shotgun sequencing. However, the ultra-deep sequencing required to profile strain level communities can be prohibitively expensive, and computationally challenging when applied to large-scale surveys of the human microbiome. Our ISR pipeline provides a high-resolution yet cost-effective approach for molecular epidemiology studies by allowing subspecies level tracking of microbiota. This method can be easily integrated into existing amplicon sequencing protocols by simply changing PCR primers, where it could provide the ability to resolve communities into a large number of variants.

This approach could be valuable for studies that address fundamental questions important for oral health. The human oral microbiome is a complex system whose composition is driven by host and environmental factors, with community alterations at the level of species being implicated in several common diseases such as dental caries and periodontitis. It is unclear if these perturbations in species abundance are simply fluctuations in levels within the existing community, or if strains adapted to the changing environment are acquired or lost. Transmission and acquisition of communities is not thoroughly understood either. Insights into these questions have far-reaching implications for therapeutic approaches. Further, the ability to accurately predict the source for a dental plaque sample analyzed using the ISR method could be useful for microbial fingerprinting and forensic tracking applications.

## CONCLUSIONS

While the 16S gene allows identification of bacteria at the level of species, it does not generally provide sufficient information to resolve strains within a species. This novel approach included two enhancements of standard 16S amplicon pipelines, a more variable target sequence, the ISR, and incorporation of a high-resolution probabilistic error modeling pipeline, DADA2. This resulted in improved resolution of communities, and revealed a highly personalized human oral microbiota.

- Specifically, the ISR-DADA2 approach detected 5.2-fold more sequence variants from dental plaque samples than the standard 16S species-level reference database approach.
- Individuals could be perfectly resolved, with no overlap seen among samples among 5 subjects over 1 year in NMDS ordinations. Two independent machine-learning based modeling approaches, hierarchical clustering and random forest classification, both converged to a perfect resolution of individuals, further validating the resolving power of the ISR approach.
- An ISR database was developed by extracting the 16S-23S intergenic spacer region from 200+ publicly available genomic sequences of common oral bacteria. Mapping to this database, multiple genotypic variants of the ISR were identified for most species, demonstrating a high level of subspecies variation in the oral microbiota. Some were widely shared and others were unique to 1 individual.

The ISR approach described here provides a high-throughput, high-resolution yet cost-effective method that allows subspecies-level community fingerprinting at a cost comparable to 16S rRNA gene amplicon sequencing. This new approach will be useful for a range of applications that require high-resolution identification of organisms, including microbial tracking, community fingerprinting and identification of virulence-associated strains.

Future development should include exploring the relationship between ISR and the rest of the genome to determine its utility as a functional marker, and to increase the scope of the ISR database to achieve comprehensive representation of the community.

## Acknowledgements

We thank Dr. Benjamin J. Callahan, from the Department of Population Health and Pathobiology, North Carolina State University, for his insights on adaptation of the DADA2 pipeline for ISR amplicon sequence processing. The research reported here was supported in part by the US National Institute of Dental and Craniofacial Research, under award numbers R01 DE024327 and R01 DE024463.

## Ethics approval and consent to participate

The studies described in this manuscript were approved by the institutional review board (IRB) of The Ohio State University (IRB study number: 2014H0176). All participants provided written informed consent.

## Availability of data and materials

The sequencing data generated during the current study were submitted to NCBI’s the Sequence Read Archive (SRA), and is accessible with SRA accession number SRP144761. The database created for the current study and all relevant meta data are available upon request.

**Table 1.** Most abundant species and the ISR-types they could be resolved into.

| Species* | ISR-types** |
| --- | --- |
| *Streptococcus mitis pneumoniae infantis oralis* | 33 |
| *Streptococcus sanguinis* | 12 |
| *Streptococcus intermedius constellatus* | 9 |
| *Haemophilus parainfluenzae* | 41 |
| *Streptococcus gordonii* | 7 |
| *Granulicatella adiacens* | 39 |
| *Leptotrichia buccalis* | 22 |
| *Gemella morbillorum* | 9 |
| *Prevotella intermedia* | 6 |
| *Gemella haemolysans* | 12 |
| *Streptococcus australis* | 5 |
| *Streptococcus cristatus* | 5 |
| *Streptococcus vestibularis salivarius* | 9 |
| *Prevotella nigrescens* | 10 |
| *Pseudomonas monteilii* | 10 |

\*In order of decreasing overall abundance. \*\*Mean = 15.3 ISR-types/species.

**Supplementary Table S1.**
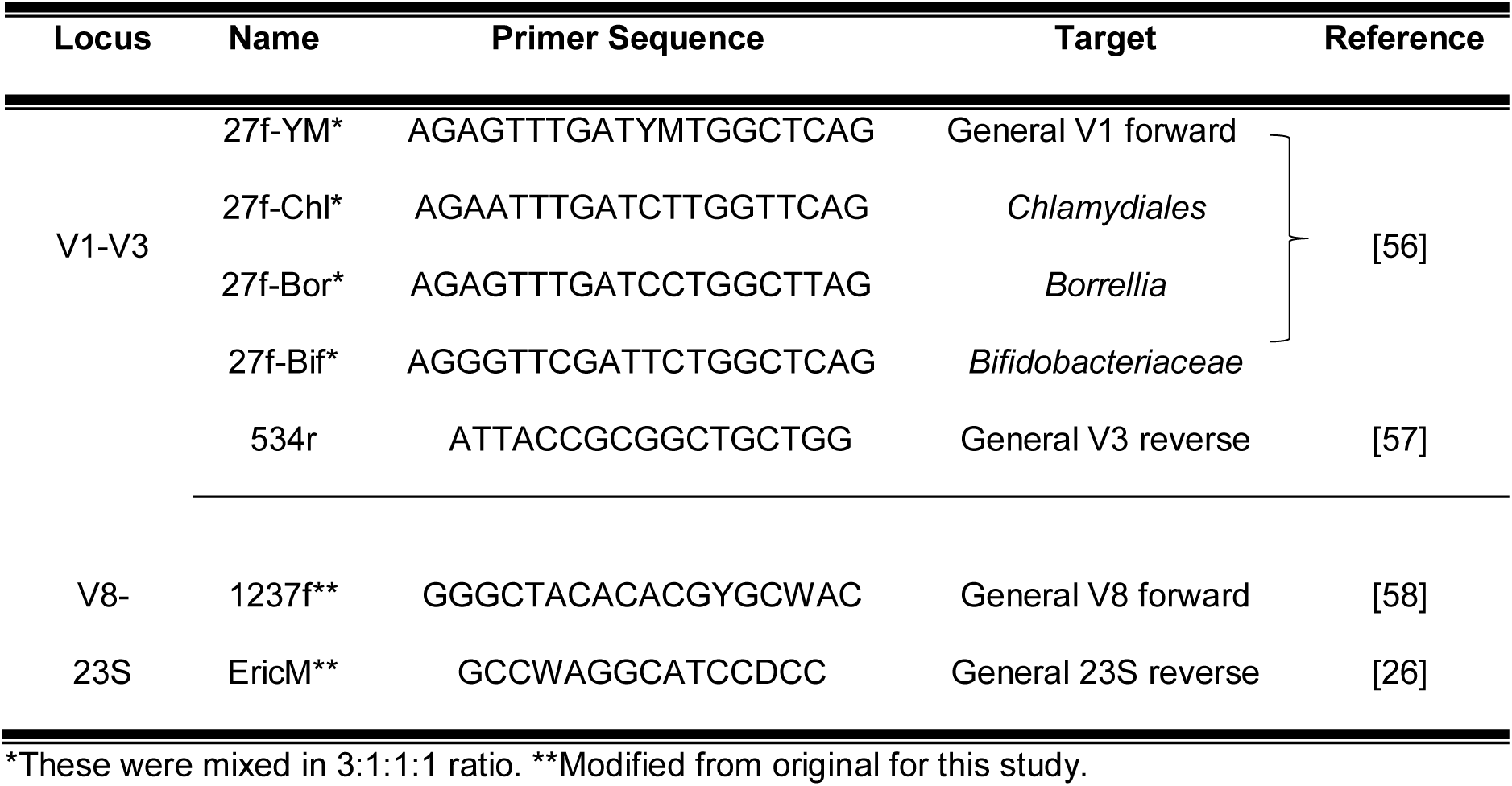
Primers used in this study.

**Figure S1.**
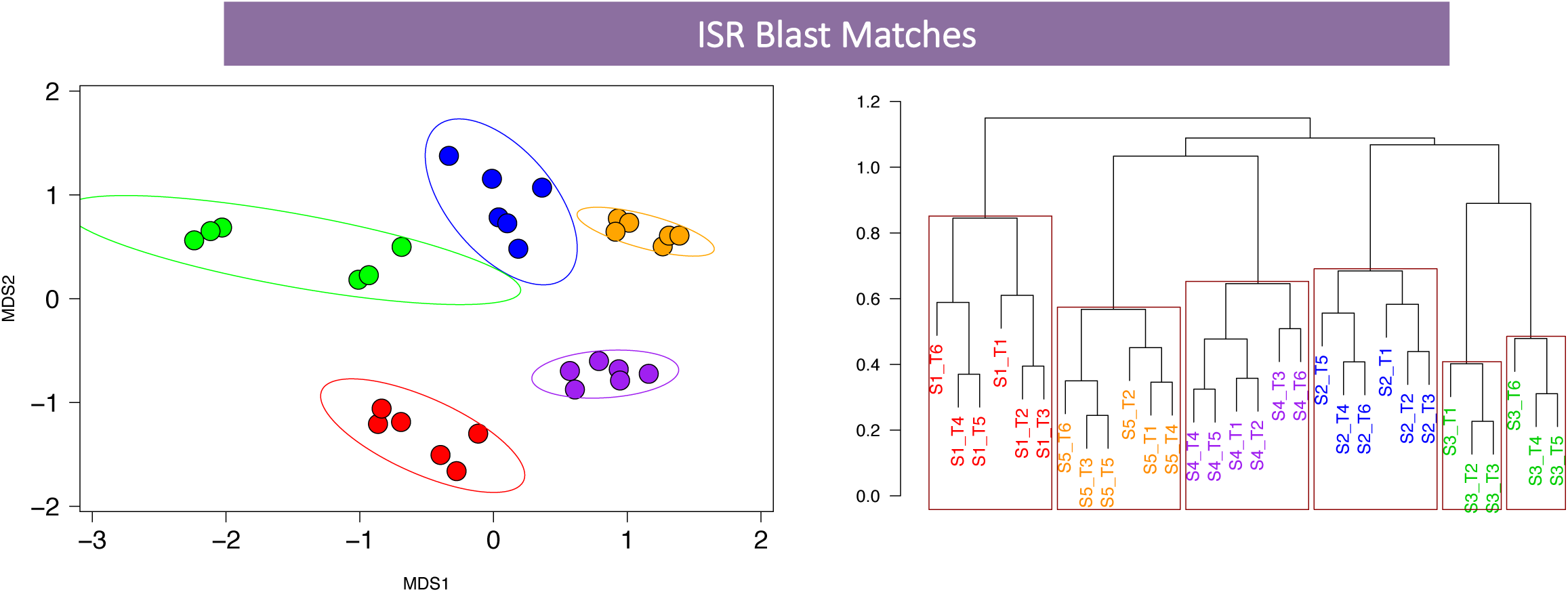
Distance based community analysis. A reduced dataset of only those ASVs that could be mapped to species still provided sufficient resolution to distinguish individuals using NMDS plots and Hierarchical Clustering Dendogram.

**Figure S2.**
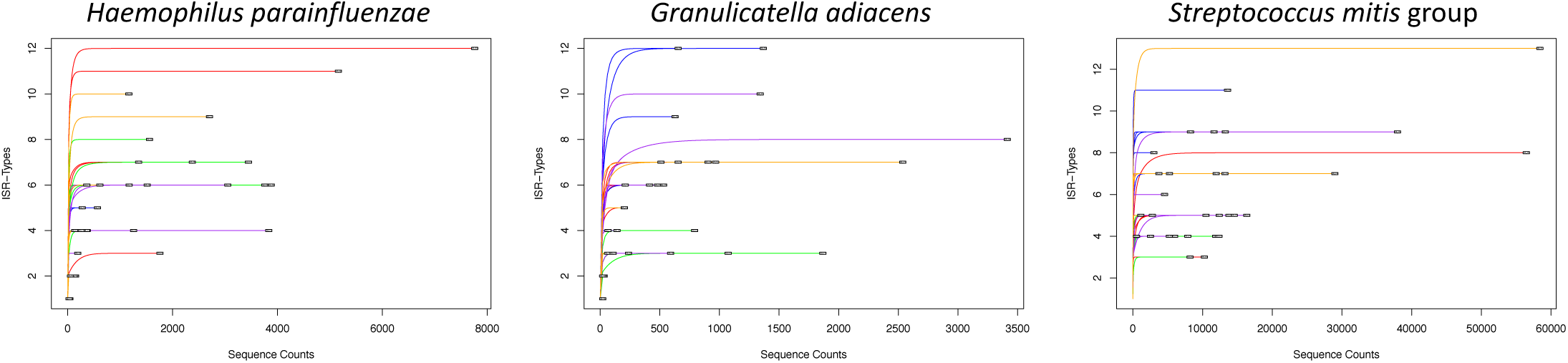
Rarefaction curve for the 3 most diverse species. For most samples, the number of observed ISR-types plateaued within the sequencing depth achieved.

